# Comparative analysis of high- and low-level deep learning approaches in microsatellite instability prediction

**DOI:** 10.1101/2022.04.09.487718

**Authors:** Jeonghyuk Park, Yul Ri Chung, Akinao Nose

**Affiliations:** Department of Physics, Graduate School of Science, The University of Tokyo, Tokyo, Japan; Pathology Center, Seegene Medical Foundation, Seoul, Korea; Department of Complexity Science and Engineering, Graduate School of Frontier Sciences, The University of Tokyo, Chiba, Japan

## Abstract

Deep learning-based approaches in histopathology can be largely divided into two categories: a high-level approach using an end-to-end model and a low-level approach using feature extractors. Although the advantages and disadvantages of both approaches are empirically well known, there exists no scientific basis for choosing a specific approach in research, and direct comparative analysis of the two approaches has rarely been performed. Using the Cancer Genomic Atlas (TCGA)-based dataset, we compared these two different approaches in microsatellite instability (MSI) prediction and analyzed morphological image features associated with MSI. Our high-level approach was based solely on EfficientNet, while our low-level approach relied on LightGBM and multiple deep learning models trained on publicly available multiclass tissue, nuclei, and gland datasets. We compared their performance and important image features. Our high-level approach showed superior performance compared to our low-level approach. In both approaches, debris, lymphocytes, and necrotic cells were revealed as important features of MSI, which is consistent with clinical knowledge. Then, during qualitative analysis, we discovered the weaknesses of our low-level approach and demonstrated that its performance can be improved by using different image features in a complementary way. We performed our study using open-access data, and we believe this study can serve as a useful basis for discovering imaging biomarkers for clinical application.

## Introduction

Deep learning techniques have demonstrated their ability to predict and extract various medical information such as diagnosis^1–5^, prognosis^6–9^ and genetic alterations^9–14^ from hematoxylin and eosin (H&E)-stained slide images. Deep learning algorithms are generally trained to predict the target of choice directly from images in a data-driven manner, and their current performances are high enough for clinical application^1, 3–5^. However, this end-to-end approach is commonly referred to as a “black box model” since it is difficult to analyze how predictions have been made^15^. Thus, to overcome this lack of interpretability, another approach is being investigated that predicts morphological features using a deep learning model and then predicts targets using these features^16, 17^. Diao et al. conducted a notable work that showed the potential of this approach^17^. They first trained a three-class (cancer tissue, cancer-associated stroma, necrosis) tissue segmentation model and a five-class (lymphocyte, plasma cell, fibroblast, macrophage, cancer cell) cell detection model to predict tissue and cells in H&E images. Then, they defined 607 image-based features from the combinations of tissue and cell information to predict molecular signatures and reported that the performance of their model was comparable to that of an end-to-end method. Furthermore, the group investigated what morphological features were related to certain molecular signatures. Such investigation has clinical significance since it can identify novel imaging biomarkers for molecular signatures and other clinical factors.

In general, end-to-end models that utilize incomprehensible high-level features (hereafter referred to as the high-level approach) outperform approaches that apply tissue- and cell-level human-interpretable image features (hereafter referred to as the low-level approach). Our first question was: What is the quantitative difference in performance between the two approaches on an identical task? High-level and low-level approaches analyze important morphological image features for target prediction in different ways^10, 17, 18^. Our second question was: When the important image features are extracted by the two approaches, are these morphological features similar? Currently, we do not have any scientific basis for selecting an approach for biomarker research. If the above two questions can be answered through direct comparison, it would be greatly beneficial for researchers in the field of pathology research since they can choose a more suitable approach for a particular study. In an attempt to answer these questions, we chose to perform our research with a microsatellite instability (MSI) task. Microsatellites are repeated 1-6 nucleotide sequences in DNA that are also called short tandem repeats or simple sequence repeats and are considered a result of DNA slippage during replication^19^. Normally, the DNA repair system called mismatch repair (MMR) corrects these errors; the loss of MMR genes in tumor cells results in microsatellite instability. MSI-high tumors tend to respond well to immunotherapy, and MSI has now become a pancancer biomarker for checkpoint inhibition therapy^20^. Currently, MSI status is generally evaluated by polymerase chain reaction assay or immunohistochemistry. However, it has been reported that MSI can be predicted from H&E images by high-level approaches^11, 12, 21^. We thought that low-level approach-based predictions would also be possible as MSI tumors are characterized by distinct morphological characteristics such as increased tumor-infiltrating lymphocytes (TIL) and poor differentiation^22^. For the dataset, the availability and wide applicability of the Cancer Genomic Atlas (TCGA)-based curated dataset for MSI prediction (hereafter, MSI dataset) of colorectal (CRC) and stomach (STAD) cohorts seemed suitable^11, 23^

In this study, we analyzed differences in performance and morphological image features extracted by high-level and low-level approaches (Figure 1a). First, we designed high- and low-level approaches and developed the models for both approaches with various hyperparameters. To prevent a performance difference due to excessive effort in one approach, the naive method was used as much as possible. For our high-level approach, we reproduced and adapted the methodology used by Kather et al.^11^. For our low-level approach and morphological feature analysis, we used three deep learning models with publicly available datasets as follows: a nine-class tissue classification model with a colorectal tissue dataset published by Kather et al.^16^; a six-class nuclei detection model with PanNuke^24^, a nuclei dataset for pancancer^25^; and a two-class gland detection model with a colon gland dataset that became available for the gland segmentation (GlaS) challenge contest^26^. The low-level approach consisted of features from these feature extractors and LightGBM (LGBM) for classifying MSI. Since the dataset trained by the tissue and gland models belongs to the colonic data, we focused on MSI prediction in the CRC cohort. Second, we compared the performance of the two approaches. Furthermore, we also compared the performance when STAD cohort data were added to the training set since the morphological characteristics of the CRC and STAD cohorts were found to be similar in our previous study^27^. Third, we extracted and compared the morphological image features related to MSI from each approach with the best-performing models. Finally, by selecting and qualitatively reviewing cases based on the concordance of the predictions of high- and low-level approaches, we uncovered the limitations of our low-level approach and a possible solution to overcome it. Our major contribution of the current study is that we demonstrated that discovering morphological characteristics related to MSI using the high-level approach is comparable to using the low-level approach and statistical data analysis. This is advantageous in that it provides a scientific basis for discovering morphological features and a biomarker through a high-level approach, as many pathology studies have adapted high-level approaches^1–6, 8–13, 18^.

**Figure 1.**
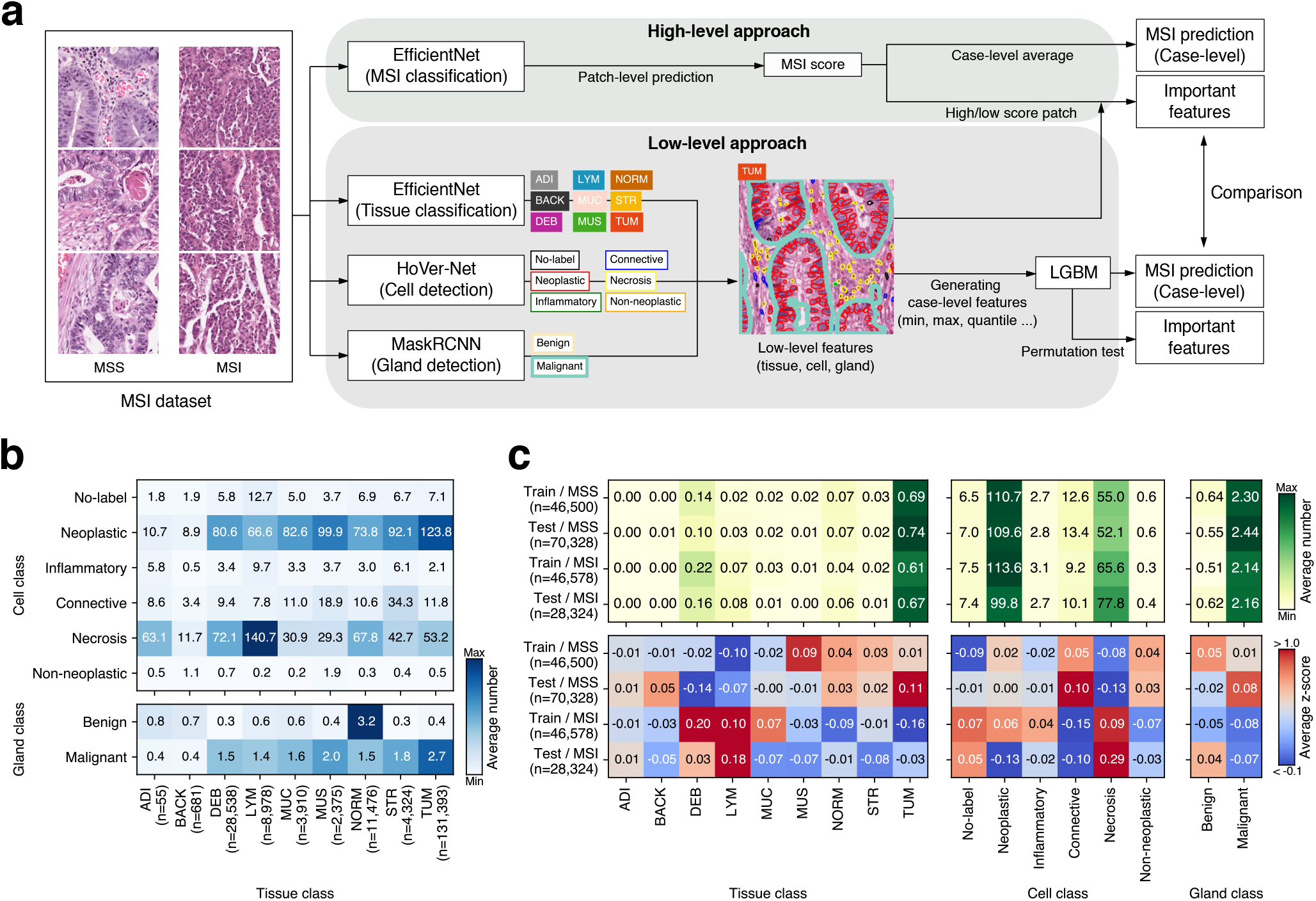
Conceptual view comparing high-/low-level approaches and dataset analysis of low-level features. (a) Pipeline for high and low-level approaches. The high-level approach uses high-level image features that utilize the deep learning architecture EfficientNet to predict microsatellite instability (MSS, microsatellite stable; MSI, microsatellite instable) directly from image patches. The low-level approach utilizes low-level features such as the number of tissues, cells, and glands. After defining and generating case-level features from low-level features, LightGBM predicts MSI from the features. (b) Correlations among low-level features derived from both the training and test sets from the colorectal cancer (CRC) cohort. (c) Low-level image features of the training and test sets of the CRC cohort. Average number (top) and z score (bottom). ADI, adipose tissue; BACK, background; DEB, debris; LYM, lymphocytes; MUC, mucus; MUS, smooth muscle; NORM, normal colon mucosa; STR, cancer-associated stroma; TUM, colorectal adenocarcinoma epithelium. The heatmaps were created with Python v3.8.8 (https://www.python.org) and Matplotlib v3.4.3 (https://www.matplotlib.org). The figure was generated using Inkscape v1.1 (https://inkscape.org).

## Results

### Deep learning models for high-/low-level approaches

We reproduced and adapted the MSI classification model using colorectal (CRC) cohort image patches in the MSI classification dataset provided by Kather et al.^11^. The methodology mainly followed their study with a few modifications. We performed fivefold cross-validation instead of defining a fixed validation set using the training set, and we defined the search space including several model architectures and hyperparameters and optimized the models (Supplementary Tables 1, 2, see Materials and Methods section for details). We used the area under the receiver operating characteristic curve (AUROC) as a model performance measure to allow comparison with previous studies^11, 18^. The model that showed the best performance in the search space was the model based on the EfficientNet architecture^28^, and the average AUROC of the fivefold models in the CRC test set was 0.8065 (95% confidence interval (CI), 0.7758-0.8373), which is higher than the AUROC of 0.76 reported by a previous study^11^ (Table 1 and Supplementary Table 3).

**Table 1.** Comparison of performance of high- and low-level approaches. Note that performances are measured only in the test set of the CRC cohort. p=0.046 for between AUROCs of high-level (CRC) and high-level (CRC+STAD), p=0.085 for between AUROCs of low-level (CRC) and low-level (CRC+STAD), p=7.4 × 10^−5^ for between AUROCs of high-level (CRC) and low-level (CRC). p values are calculated from Student’s t test. The critical significance level adjusted by Bonferroni correction is p=0.025. AUROC, area under the receiver operating characteristic curve; CI, confidence interval; LGBM, light gradient boosting machine; CRC, colorectal; STAD, stomach.

| Approach | Training cohort | AUROC (95% CI) |
| --- | --- | --- |
| High-level (EfficientNet) | CRC | 0.8065 (0.7758-0.8373) |
|  | CRC+STAD | 0.7678 (0.7341-0.8015) |
| Low-level (LGBM) | CRC | 0.6749 (0.6365-0.7134) |
|  | CRC+STAD | 0.6445 (0.6250-0.6640) |

Next, we prepared three deep learning models for morphological feature extraction. A tissue classification model was optimized similarly to the MSI classification model using the colon tissue classification dataset published by Kather et al. which contains nine tissue classes as follows: adipose tissue (ADI), background (BACK), debris (DEB), lymphocytes (LYM), mucus (MUC), smooth muscle (MUS), normal colon mucosa (NORM), cancer-associated stroma (STR), colorectal adenocarcinoma epithelium (TUM)^16^. Our best model an achieved average overall accuracy of 0.9523 (95% CI, 0.9424-0.9623) on the test set of the tissue dataset which is higher than the 0.943 of a previous study^16^ (see Supplementary Table 4 for optimization of this model and different metrics). The tissue classification dataset was extracted from CRC tissue, which is the same as the MSI classification dataset, and a model trained on this dataset was used in the CRC cohort of the TCGA dataset^16^.

Then, we obtained Hover-Net24 pretrained with the PanNuke dataset^25^ as a nuclei detection model. This model was expected to work well with the MSI dataset since the PanNuke dataset includes the COAD (colon adenocarcinoma) cohort of the TCGA dataset, and HoVer-Net was used to classify nuclei in the COAD and READ (rectal adenocarcinoma) cohorts of the TCGA dataset^18, 25^. The model allowed us to detect nuclei into 6 classes: no-label, neoplastic, inflammatory, connective, necrosis, and non-neoplastic nuclei.

Finally, we trained a MaskRCNN^29^ for two-class gland detection using the GlaS contest dataset^26^. The GlaS contest data consist of colonic tissue data, including TCGA-COAD, which we expected to work on the MSI dataset as well. We achieved a competent performance of our gland detection model (single-class F1-score of 0.9019 in test set A and 0.7290 in test set B) which corresponds to the second place in the GlaS challenge. Before proceeding to the next analysis, one pathologist (Y.R.C.) visually reviewed the results of the three models to examine whether they were qualitatively acceptable.

### Image feature characteristics of the datasets

Certain classes of low-level image features were expected to have correlations with one another. For example, in a TUM patch, since it includes neoplastic cells and malignant glands, it would yield a high number of neoplastic cells and malignant glands. A normal tissue patch would have a high number of benign glands, and an STR patch would have many connective tissue cells. We compared the number of cells (equivalent to the number of nuclei) and glands based on the tissue classification results in the CRC cohort of the MSI dataset (Figure 1b). As expected, correlations between TUM-neoplastic cell and NORM-benign gland were shown, and many connective cells were also observed in the stroma and muscle. On the other hand, in the case of the LYM patch, there were more necrotic cells than inflammatory cells, and we believe this may be due to the similarity of the small, hyperchromatic nuclei of necrotic and inflammatory cells, which may be difficult to accurately distinguish at the nuclear level. Overall, the high number of neoplastic and malignant glands may have resulted from the fact that the MSI dataset itself was sampled from the tumor region^11^. Next, we investigated the image feature characteristics for MSS (microsatellite stable)/MSI (microsatellite instable) inherent in the dataset (Figure 1c). MSS showed a high z score in NORM, STR, TUM, connective, neoplastic, and malignant glands in both the training and test sets. In contrast, high z scores for DEB, LYM, and necrosis were observed in MSI, which is in line with clinical findings of high TIL infiltration and poor differentiation^22^.

### Image features related to microsatellite instability extracted by high- and low-level approaches

To analyze the morphological features according to the MSI score yielded by the high-level approach, average statistics of the image features were calculated in the patches in the top and bottom 50% and 10% MSI scores of the CRC test set. Overall, image features extracted from the entire image patch and each MSS/MSI patch appeared similar to the trends inherent in the dataset (Figure 2a). The counts of of NORM, TUM, neoplastic, connective, non-neoplastic, and benign/malignant glands were high in MSS, and the counts of DEB, LYM, and necrosis were high in MSI (Figure 3). For the low-level approach, we aggregated patch-level image features into case-level features using simple feature engineering. For each patch, the probabilities of tissue classes and the number of cells and glands were converted to case-level by the following function: average, maximum, minimum, standard deviation, and 25, 50, and 75 quantiles. Through this process, the 17 patch-level image features became 119 case-level image features. Using this case-level feature on the CRC training set, LightGBM (LGBM) was optimized as an MSI classifier and showed an average AUROC of 0.6749 (95% CI, 0.6365-0.7134) on the CRC test set, which was significantly lower than that of the high-level approach using the end-to-end model (Table 1). To analyze features related to MSI prediction, we performed a permutation test to acquire feature importance by measuring the performance difference with each image feature. As a result, among the features that showed that the upper boundary of the 95% CI of permutated performance was lower than its original value, we found that DEB, LYM, NORM, STR, necrosis, and malignant gland were notable features that affected performance (Figure 2b). DEB, LYM, and necrosis appeared to be important features in both high and low-level approaches, suggesting that they were important factors in predicting MSI. Next, we conducted cohort aggregation by adding stomach (STAD) cohort data to the training set in both approaches. We expected an increased performance since normal colonic and gastric tissues share similar characteristics^27^. However, adding stomach (STAD) cohort data did not improve the performance of either approach (Table 1), implying that the image features of MSI of CRC and STAD may be different. Further analysis confirmed that the image feature characteristics of STAD data differed from CRC data as different trends were observed for DEB, LYM, and necrosis (Supplementary Figure 1).

**Figure 2.**
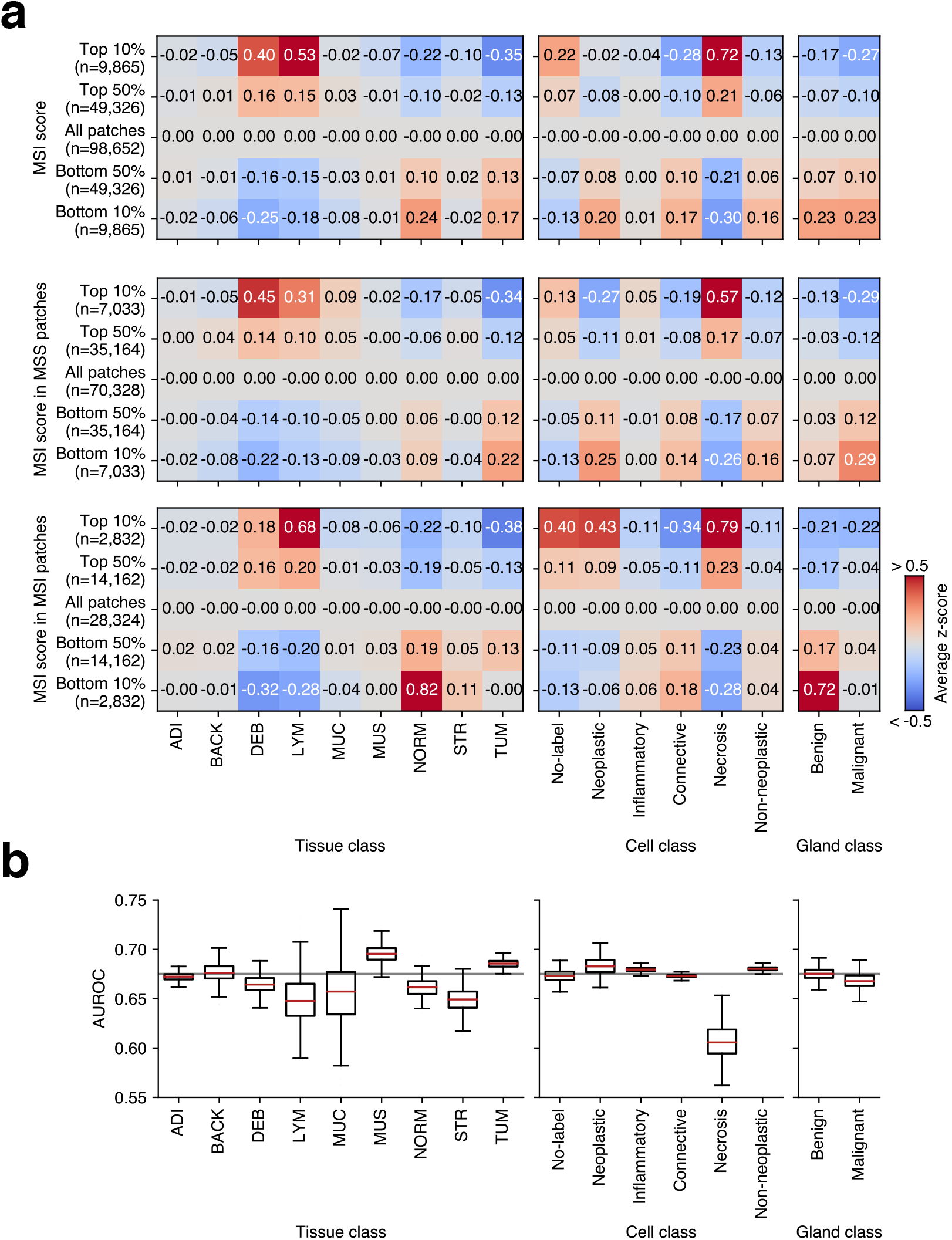
Image features extracted by high- and low-level approaches. (a) For the high-level approach, average z scores for tissue, cell, and gland features were displayed as a heatmap by dividing patches into five groups (top 10, 50%, all patches and bottom 10, 50%) based on the MSI score. The values are calculated in all patches (top), and in MSS patches (middle), in MSI patches (bottom). (b) For the low-level approach, the values of the areas under the receiver operating characteristic curve (AUROCs) corresponding to each feature calculated from the permutation test are shown as boxplots. The gray line indicates the original performance (0.6749) of the model. ADI, adipose tissue; BACK, background; DEB, debris; LYM, lymphocytes; MUC, mucus; MUS, smooth muscle; NORM, normal colon mucosa; STR, cancer-associated stroma; TUM, colorectal adenocarcinoma epithelium. The heatmaps were created with Python v3.8.8 (https://www.python.org) and Matplotlib v3.4.3 (https://www.matplotlib.org). The figure was generated using Inkscape v1.1 (https://inkscape.org).

**Figure 3.**
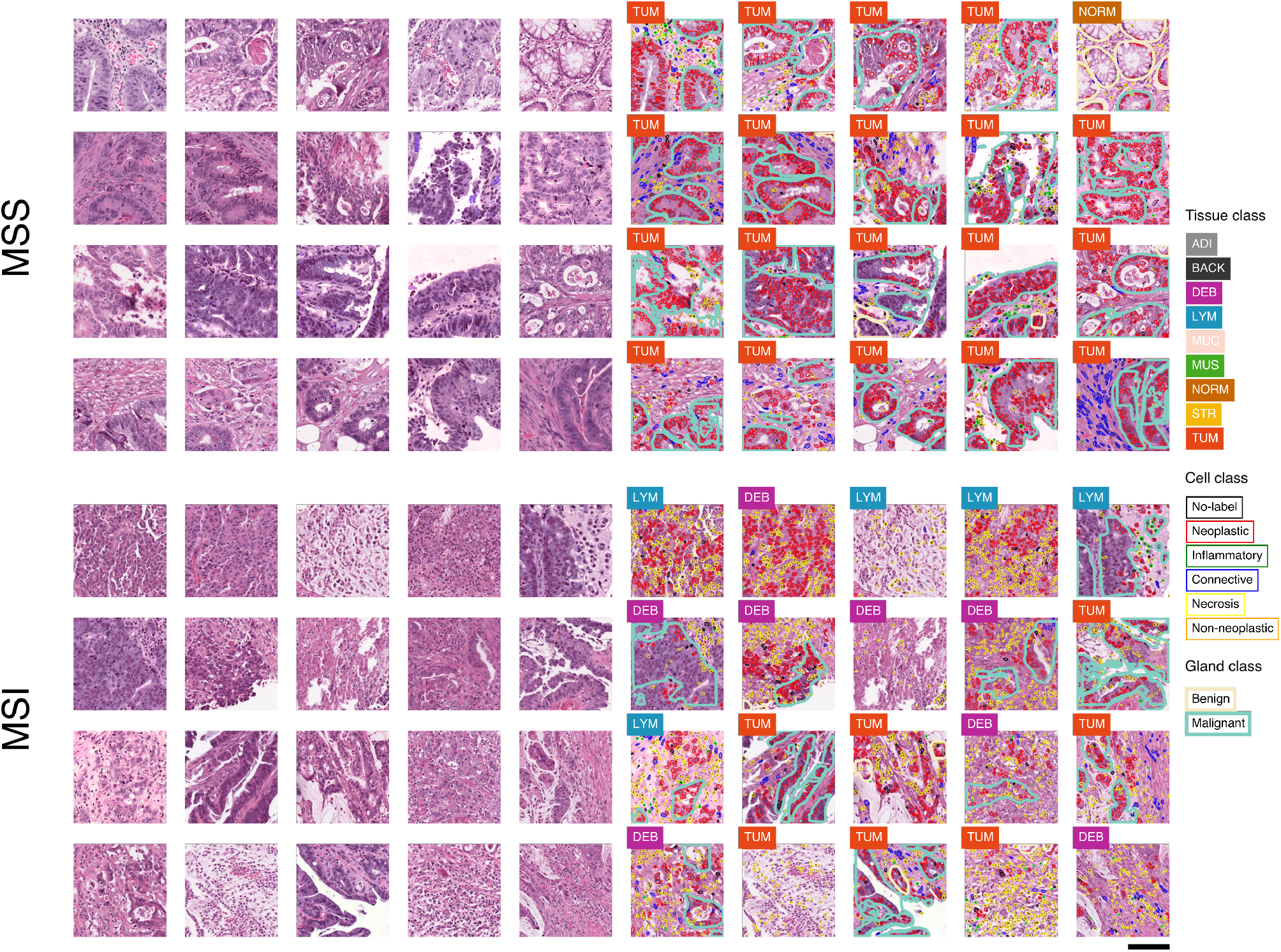
Examples of H&E image patches and predictions. Unprocessed patches from the test set of the colorectal (CRC) cohort alongside corresponding visualizations of tissue, nuclei, and gland predictions. The 20 highest and lowest MSI score patches, which do not overlap at the case-level, are shown as MSS (top) and MSI (bottom). MSS, microsatellite stable; MSI, microsatellite unstable (instability); ADI, adipose tissue; BACK, background; DEB, debris; LYM, lymphocytes; MUC, mucus; MUS, smooth muscle; NORM, normal colon mucosa; STR, cancer-associated stroma; TUM, colorectal adenocarcinoma epithelium. Scale bar, 50 *μ*m. The figure was generated using Inkscape v1.1 (https://inkscape.org).

### Qualitative review of select cases

The large performance gap between the high-level approach and the low-level approach suggests that undefined image features relevant to MSI prediction exist. However, it is almost impossible to find such features while qualitatively reviewing hundreds of thousands of patches. Thus, we decided to look for these potential features by qualitatively reviewing high-score image patches from the cases that the high-level approach correctly predicted and the low-level approach did not (Figure 4). We selected six MSS cases and six MSI cases based on the difference between high-level and low-level predictions, and in half of them, our low-level approach made incorrect predictions (Supplementary Figure 2). The unprocessed H&E patches and their processed results were reviewed while recording the following histologic characteristics for each case: tumor differentiation (well, moderate, poor), tumor cluster morphology (tubular, papillary, oval, solid, cribriform, trabecular, micropapillary, etc.), and tumor cell morphology, including both nuclear (pleomorphic, hyperchromatic, vesicular) and cytoplasmic features (shape, amount, color) (Table 2). Two of the three MSS cases with incorrect prediction by the low-level approach showed abundant vesicular nuclei; two of the three MSI cases with incorrect prediction by the low-level approach showed a predominant tubular/papillary structure of the tumor glands (Table 2 and Figure 5a, b). Unfortunately, we were not able to quantitatively confirm the importance of vesicular nuclei and tubular/papillary structure since it was difficult to design such features using the low-level models we used. Next, we observed that the patches classified as LYM (lymphocyte) tissue generally resulted in a large number of necrotic cell nuclei (Figure 5c) which should have been classified as inflammatory cell nuclei. Therefore, we supposed that the number of lymphocytes (inflammatory cell nuclei) may be an important feature that was omitted. We trained LGBM by adding the number of necrotic cell nuclei in the LYM patch as a new feature and confirmed that the performance was improved from the average AUROC of 0.6749 to 0.6785. This result suggests that the number of lymphocytes is an important factor in MSI prediction.

**Figure 4.**
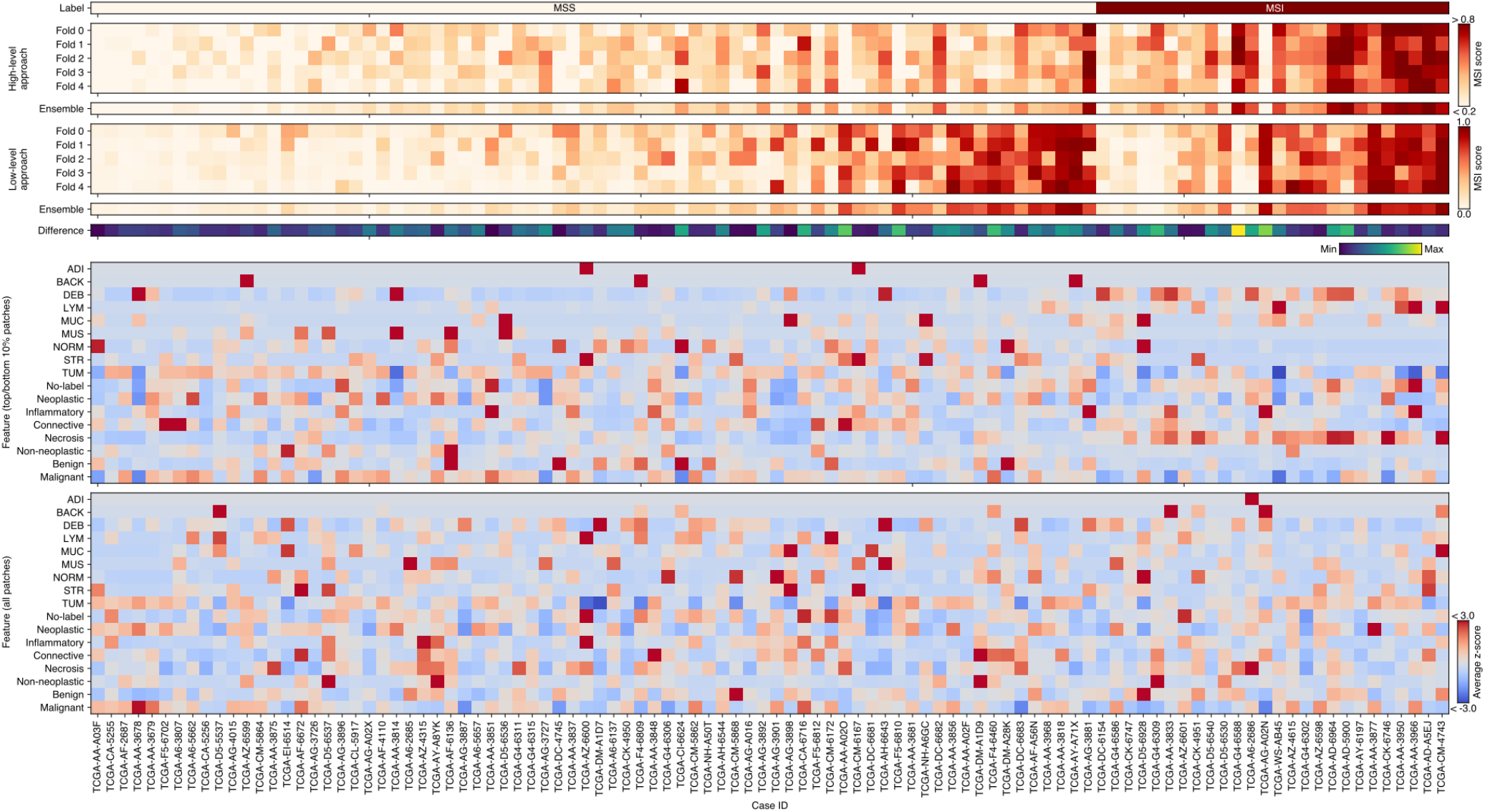
Comparison of case-level prediction of high and low-level approaches. Case-level prediction of EfficientNet (high-level approach) and LGBM (low-level approach) with MSI labels (top six panels). The predicted MSI scores of fivefold models and their ensemble are shown. Their difference is calculated based on the z scores of each ensemble. Representative image features derived from high/low-level approaches (bottom two panels). The average z score of the number of tissues, cells, and glands of the top 10% of MSS/MSI patches and the case-level average. TCGA case IDs are shown as column labels. LGBM, light gradient boosting machine; MSS, microsatellite stable; MSI, microsatellite instable (instability); TCGA, the cancer genome atlas; ADI, adipose tissue; BACK, background; DEB, debris; LYM, lymphocytes; MUC, mucus; MUS, smooth muscle; NORM, normal colon mucosa; STR, cancer-associated stroma; TUM, colorectal adenocarcinoma epithelium. The heatmaps were created with Python v3.8.8 (https://www.python.org) and Matplotlib v3.4.3 (https://www.matplotlib.org). The figure was generated using Inkscape v1.1 (https://inkscape.org).

**Table 2.** Qualitative review of select cases. HA and LA indicate the scores of high and low-level approaches, respectively. A high score corresponds to MSI (=1). See Supplementary Figure 2 for additional details for select cases. MSS, microsatellite stable; MSI, microsatellite instable.

| Case ID | Label | HA | LA | Tumor differentiation | Tumor cluster morphology | Tumor cell morphology |
| --- | --- | --- | --- | --- | --- | --- |
| TCGA-AA-A03F | MSS | 0.05 | 0.02 | well | tubular | columnar |
| TCGA-AF-2687 | MSS | 0.18 | 0.01 | moderate | irregular | pleomorphic |
| TCGA-CA-5255 | MSS | 0.15 | 0.04 | moderate | irregular, cribriform | pleomorphic, ample eosinophilic cytoplasm |
| TCGA-AA-A02O | MSS | 0.10 | 0.7 | well | tubular | columnar |
| TCGA-F4-6460 | MSS | 0.24 | 0.82 | moderate | oval | vesicular, slightly columnar |
| TCGA-F5-6810 | MSS | 0.15 | 0.72 | moderate | irregular, cribriform | vesicular, slightly columnar |
| TCGA-A6-2686 | MSI | 0.60 | 0.21 | poor | solid | pleomorphic |
| TCGA-AD-5900 | MSI | 0.78 | 0.37 | moderate | tubulopapillary | columnar |
| TCGA-G4-6588 | MSI | 0.73 | 0.04 | moderate | tubulopapillary | slightly columnar |
| TCGA-AA-3966 | MSI | 0.76 | 0.83 | poor | single cell, diffuse, solid, cord-like | dark basophilic |
| TCGA-AD-A5EJ | MSI | 0.81 | 0.79 | moderate | irregular, tubulopapillary, micropapillary | columnar |
| TCGA-CM-4743 | MSI | 0.72 | 0.97 | poor | solid, diffuse, micropapillary | pleomorphic |

**Figure 5.**
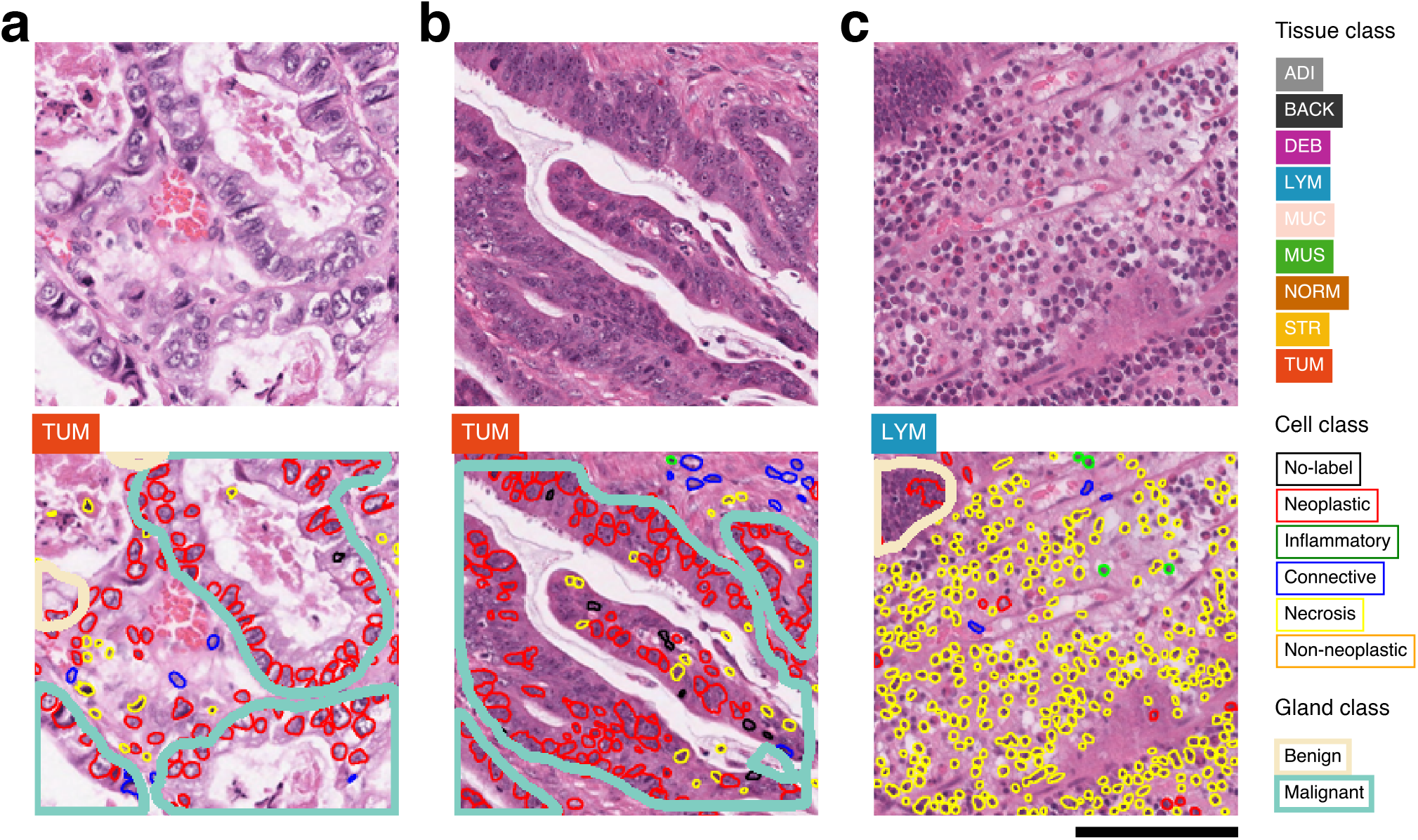
Representative patch samples. Patches sampled from TCGA-F5-6810 (a), TCGA-G4-6588 (b), and TCGA-AD-5900 (c). (a) Vesicular cells (b) A predominant tubular/papillary structure of the tumor glands is shown. (c) Patches classified as LYM tissue. Inflammatory cells (lymphocytes) are misclassified as necrotic cells. ADI, adipose tissue; BACK, background; DEB, debris; LYM, lymphocytes; MUC, mucus; MUS, smooth muscle; NORM, normal colon mucosa; STR, cancer-associated stroma; TUM, colorectal adenocarcinoma epithelium. Scale bar, 50 *μ*m. The figure was generated using Inkscape v1.1 (https://inkscape.org).

## Discussion

Generally, end-to-end models (high-level approach) are data-efficient and can achieve a high performance relatively easily compared to the low-level approach. In our study, the low-level approach showed a significantly lower performance than the high-level approach. However, with sufficient feature engineering, the performance of the low-level approach can be equivalent to that of the high-level approach, as shown in the study by Diao et al.^17^. Such feature engineering (which includes complex features such as the density of lymphocytes within 80 *μ*m of the cancer–stroma interface) is difficult, which makes it a common disadvantage of the low-level approach. Thus, the low performance of our low-level approach suggests that several features related to MSI may exist in addition to the features we defined. To find these features, we selected cases using both high- and low-level approaches and conducted a qualitative review. A qualitative review of select cases suggested that vesicular nuclei and tubular/papillary structure of tumor glands may be potential morphological features of classification of MSS/MSI (Table 2). These features were not involved in our low-level approach but were likely incorporated as image features in the high-level approach, which may be one of the reasons for the higher performance of our high-level approach. Additionally, during histological review, we observed that many of the lymphocytes in LYM patches were detected as necrotic cells (Figure 5c). This is also shown indirectly in the correlation analysis among image features (Figure 1b). After applying the necrotic cells in LYM patches as a new feature to the low-level approach, performance was improved. This finding suggests that the tissue classification model can be used in a complementary way to enhance performance since nuclei classification often relies on the spatial context of the surrounding tissue.

For a high-level approach, adapting weakly supervised learning or multiple instance learning should be conducted to identify morphological features more specific to MSI since the morphological features of MSS or MSI can appear locally^1, 18^. Bilal et al.18 presented a deep learning framework (which can be classified as a high-level approach) using weakly supervised learning and reported that it achieved an AUROC of 0.90 on the MSI prediction task. Additionally, they applied PanNuke pretrained HoVer-Net to top-scored image tiles and derived cell features related to MSI using feature importance calculated from a support vector machine. They reported that a high proportion of inflammatory and necrotic cells and a low proportion of neoplastic and connective cells were associated with MSI. Considering the possibility of misclassification of inflammatory cells from this study, this is in line with our results. Even though the same cell detection model (HoVer-Net with PanNuke) was used, the difference in inflammatory cells was presumed to be due to data. We used color normalized presampled images by Kather et al.^11^, while Bilal et al. used images sampled using their own method. This suggests that the pretrained HoVer-Net may provide different results depending on the image quality, which should be evaluated in future studies.

In our analysis of morphological features associated with MSI in high- and low-level approaches, we observed that DEB, LYM, and necrosis were important features for MSI in both approaches (Figure 2). Of note, this is also characteristic of the dataset itself (Figure 1c). These results suggest that a morphological feature analysis using patches selected from a high-level approach is a muefficient method since it does not require feature extraction for the entire image patch. Considering that many conventional high-level approach studies have qualitatively reviewed high-score patches related to their predictive task^1, 7, 9, 11, 13, 18^, this would have an impact in that the patches can be quantitatively analyzed on a scientific basis. For example, as Fu et al.9 extracted patches related to a mutation or prognosis in various organs, one may be able to find related biomarkers by applying our low-level feature models to the patches.

We observed that adding the STAD dataset to the CRC dataset did not result in performance enhancement of our approaches (Table 1). A possible explanation may be that morphological features related to MSI are different for the colon and stomach. In colon cancer, an increased number of tumor infiltrating lymphocytes, poor (medullary) differentiation, mucinous differentiation, and a Crohn-like reaction are considered morphological features associated with MSI-high tumors^30^. For gastric cancer, Mathiak et al. evaluated phenotypic features characteristic of MSI gastric cancers in a small case series and observed that MSI gastric tumors more frequently showed highly pleomorphic cells with large vesicular nuclei in a trabecular, nested, microalveolar, or solid growth pattern as well as inflammatory cell-rich stroma; however, these findings need further validation^31^. Thus, while the phenotypic features associated with MSI-high colorectal cancer seem relatively consistent, those for MSI-high gastric cancer remain to be elucidated.

This study has limitations. Since our study is based on patches sampled from a tumor area, the study has a bias where it inevitably results in many TUM patches with tumor-related cells and glands. Therefore, important low-level morphological features that may exist in the nontumor regions may have been missed. Additionally, since the patches consist of images sampled at a specific magnification (112 *μ*m x 112 *μ*m per patch), morphological features pertaining to wider areas (such as tumor cluster size) are not considered. Another limitation is that our low-level model was trained only with simple features, i.e., the number of targets. Adding image features such as the size and color of cells and glands may improve low-level model performance. There was also a limitation in cell and gland detection in that it was difficult to detect heterogeneous cells or glands. In particular, in malignant glands, there were cases where it was difficult to distinguish individual glands due to their markedly atypical shape. Furthermore, our gland detection model was trained on a relatively small amount of data, and the resultant low gland detection capability made it difficult to assess the impact of gland features on MSI.

We were motivated by the lack of methodological guidelines or scientific bases on which to analyze morphological features for a specific prediction task in pathology. We systematically compared performances and extracted morphological features of high-level and low-level approaches for the first time. We showed similarities and differences between the two methods and demonstrated that low-level models can be constructed in combination to improve performance. We also demonstrated that high/low-level approaches and statistical analysis of the dataset itself are both important in analyzing image features, suggesting that they can be selected and used according to the scale of a dataset or the purpose of study. We utilized open sources from datasets to models, and we have disclosed the models and codes used in this study. We believe that the current study can serve as a foundation for developing deep learning approaches in pathology for medical applications.

## Materials and Methods

### Dataset

We used three publicly available datasets in this study: histological images for MSI vs. MSS classification in gastrointestinal cancer, FFPE samples (MSI dataset)^11^; 100,000 histological images of human colorectal cancer and healthy tissue (Tissue dataset)^16^; and the GlaS challenge contest dataset (GlaS dataset)^26, 32^. They were used for the MSI classification task, tissue classification, and gland detection. The links for downloading are provided in the data availability section. Since the training set and test set are already defined in all of these datasets, five folds were generated in the training set for model training. For the MSI dataset, folds were split at the case level (patient level) so that the number of cases in each fold was similar (see Supplementary Table 2 for fivefold configuration details). We also defined a fivefold configuration for the Tissue dataset and GlaS dataset. The folds were generated uniformly at the patch level since no case-level information was included. The image resolution was 0.5 *μ*m for both the MSI dataset and the tissue dataset and 0.62 *μ*m for the GlaS dataset. All experiments were conducted on open-access data and were performed in accordance with relevant guidelines and regulations.

### High-level MSI classification model

For training of the MSI classification model, we largely adapted Kather et al.’s hyperparameters and defined a search space for finding a more optimized model^11^ (see Supplementary Table 1 for comparison). Four model architectures were included in the search space: ResNet^18^ used in Kather et al.^11, 33^; ShuffleNet used in papers that were recently published by their research group on MSI classification^12, 21, 34, 35^; and ResNext50 and EfficientNet, which are widely used and well-known for showing state-of-the-art performance^28, 36^. All models were trained from ImageNet pretrained weights and were trained in two ways such that the weights were partially frozen or fully trainable (see Supplementary Table 3 for the number of trainable parameters). We used the Adam optimizer with L2-regularization of 10^−4^ for training and we tested learning rates of 10^−3^, 10^−4^, 10^−5^ and 10^−6^. The batch size was defined for 12 GB GPU memory: 84 for ResNext, 96 for EfficientNet, and 256 for ShuffleNet and ResNet. generated so that the numbers of MSS and MSI patches were equal. Binary cross-entropy loss is used as the target loss function. Augmentation was applied with a 50% probability of vertical/horizontal flipping and affine transformation of 5-pixel shearing. One epoch was defined as 256 iterations (training steps). At the end of each epoch, accuracy was measured in the validation fold. Training was halted if the validation accuracy in a validation fold did not increase for six successive validation checks or the number of training epochs reached 100. The performance (AUROC) of the optimized models in each search space was 0.7991 (95% CI, 0.7720-0.8261) for ResNet18, 0.7544 (95% CI, 0.7039-0.8049) for ShuffleNet, 0.7871 (95% CI, 0.7790-0.7953) for ResNext50, and 0.8065 (95% CI, 0.7758-0.8373) for EfficientNet. The hyperparameters of the best EfficientNet model were a batch size of 96, a learning rate of 10^−5^ and partially frozen weights (41 trainable tensors and 1,895,698 trainable parameters). The detailed results of MSI classification model optimization are summarized in Supplementary Table 3.

### Tissue classification model

Our tissue segmentation model was optimized using the EfficientNet model architecture, an architecture with the highest performance in the MSI classification model. We used the Adam optimizer with L2 regularization of 10^−4^ for training, and we tested learning rates of 10^−3^, 10^−4^, 10^−5^, and 10^−6^. Models were trained from ImageNet pretrained weights with partially frozen or fully trainable conditions. We used balanced sampling so that the model sees each class of data equally. The batch size, loss, augmentation, definition of epoch, and other conditions were the same as those used in the optimization of the MSI classification model. The optimized model was archived with the following hyperparameters: a batch size of 96 and learning rate of 10^−5^ with fully trainable weights. It showed an average accuracy of 0.9523 (95% CI, 0.9424-0.9623) in the test set of the tissue classification dataset.16 The parameters used for tissue segmentation model optimization are summarized in Supplementary Table 4.

### Cell (nuclei) detection model

We used PanNuke pretrained HoVer-Net for cell detection^24, 25, 37^. HoVer-Net showed a high level of performance in the PanNuke dataset benchmark^37^. For inference, patches of the MSI dataset were upsampled from 0.5 *μ*m (224px x 224px, equivalent to 20x) to 0.25 *μ*m (448 px x 448 px, equivalent to 40x) using bicubic interpolation.

### Gland detection model

MaskRCNN was trained using the GlaS dataset for gland detection. The dataset contains 165 H&E images with benign and malignant gland segmentation masks (annotations). The images and corresponding annotations of the GlaS dataset were upsampled from 0.62 *μ*m to 0.5 *μ*m using bilinear and nearest interpolation, respectively. We optimized the model at learning rates of 10^−3^, 10^−4^, and 10^−5^ with hyperparameters as follows: maximum epochs=200, momentum=0.9, weight decay=0.0005. Vertical flip and horizontal flipping with 50% probability were applied as augmentation and non maximum suppression was applied to the segmented objects. A segmented object showing intersection over union (IoU) over 0.5 with a ground truth object was counted as a true positive; otherwise, it was counted as a false-positive. A false negative is defined as a ground truth object that has no corresponding detected object or shows less than an IOU of 0.5 by its corresponding detected object. The F1-score of the validation fold was calculated every 10 epochs (see the Metrics subsection for the definition of the F1-score). In the cross-validation process, the highest F1-score was achieved at a learning rate of 0.001, and the average number of validation steps was 11. Based on this cross-validation training parameter, we trained a single model with a learning rate of 0.001 with 110 epochs. The results of the gland detection model are summarized in Supplementary Table 5.

### Low-level MSI classification model

We generated case-level features to train LGBM to predict MSI at the case level. The probabilities of tissue classes and the number of cells (=nuclei) and glands were grouped by case level using these seven functions: minimum (min), maximum (max), average (avg), standard deviation (std), and 25%, 50%, and 75% quantiles. Through this case-level feature generation, the 17-dimensional features at the patch level became 119-dimensional features at the case level. The input features are scaled the z score, which is defined as *z* = (*x* - (*mean of x in train folds*) / (*standard deviation of x in train folds*). The fivefold configuration was the same as the high-level approach at the case level, and the model was searched in the following search space: 0.1, 0.01, and 0.001 for learning rate; 1.0, 0.8, and 0.5 for both bagging fraction and feature fraction; 0.0, 0.1, and 0.2 for lambda l1 and l2. Other hyperparameters were metric=binary, number of trees=31, iterations=5000, early stopping=300, and boosting type=gradient boosted decision trees. We selected the model showing the highest performance on the test set. To calculate feature importance, a permutation test was conducted. Case-level features derived from each low-level feature were permutated randomly 500 times, and the permutated performances were evaluated. For example, min, max, avg, std, and 25%, 50%, and 75% quantiles of counts of necrosis cells were permutated to archive the feature importance of counts of necrosis cells. The optimization result of the low-level approach is summarized in Supplementary Table 6.

### Metrics

AUROC is defined as the area under the *sensitivity*-(1 – *specificity*) curve. Other metrics used in this study are defined as follows:

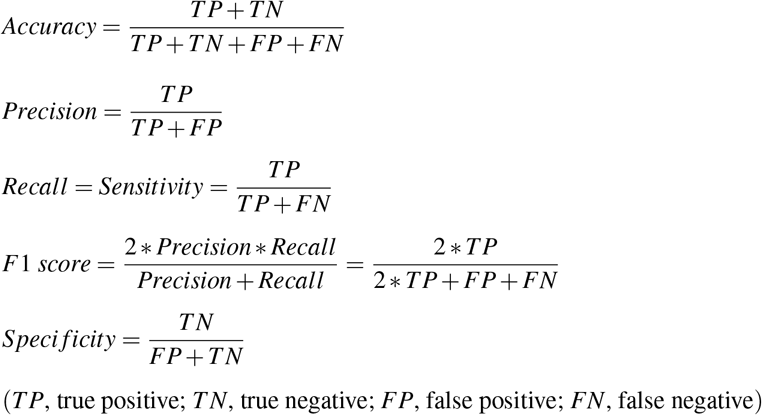

The performance evaluation of the high- and low-level approaches using the metrics is summarized in Supplementary Figure 3.

### Cohort aggregation

We added the STAD cohort data to the MSI dataset for training data. For the high-level approach, EfficientNet was trained on both the CRC and STAD cohort training sets with the hyperparameters of the best model trained on the CRC cohort training set (Supplementary Table 2). We used the same hyperparameters of the best EfficientNet model: batch size of 96, learning rate of 10^−5^, and partially frozen weights (41 trainable tensors, 1,895,698 tranable parameters). For the low-level approach, LGBM was optimized in the same search space used in optimization of the low-level approach: 0.1, 0.01, and 0.001 for the learning rate; 1.0, 0.8, and 0.5 for both bagging fraction and feature fraction; 0.0, 0.1, and 0.2 for lambda l1 and l2 using both of CRC and STAD cohort training sets.

### Case review

The review was conducted by Y.R.C, a pathologist with 5 years of posttraining experience. Based on the difference in case-level prediction, we selected 12 cases (Table 2 and Supplementary Figure 2). In these cases, we sampled the patches having the top 10% and bottom 10% of the MSI scores of the high-level approach for review.

### Environment

All of the analyses except the training and inference of MaskRCNN (gland detection model) were performed on Python version 3.8.8 and PyTorch version 1.8.1 with CUDA 11.1 and cuDNN 8 (base Docker image tag: pytorch/pytorch: 1.8.1-cuda11.1-cudnn8-runtime). For the training and inference of MaskRCNN, we used Python version 3.7.11 and PyTorch version 1.11.0 with CUDA 11.3 and cuDNN 8 (base Docker image tag: pytorch/pytorch:1.10.0-cuda11.3-cudnn8-runtime). We used the lightgbm python package version 3.3.1 for the LGBM. Additional details about the environmental configuration were provided along with our published code (see Code Availability section for the link).

## Code Availability

Source codes with model weights for the test are available at https://github.com/jeonghyukpark/msi_highlow.

## Data Availability

Image patches for MSI classification task are available at https://doi.org/10.5281/zenodo.2530835. We used only patches of formalin-fixed paraffin-embedded (FFPE) diagnostic slides and followed their train/test set split. Tissue classification data is available at https://doi.org/10.5281/zenodo.1214456. The gland segmentation (GlaS) challenge dataset is available at https://warwick.ac.uk/fac/cross_fac/tia/data/glascontest/. Code for HoverNet model and PanNuke trained weights are available at https://github.com/vqdang/hover_net.

## Author contributions

J.P. conceived the experiment(s). J.P., Y.R.C. conducted the experiment(s). J.P., Y.R.C., A.N. analyzed the results. J.P., Y.R.C., A.N. wrote the manuscript. All authors reviewed the manuscript.

