## Supplementary Information for "Comparative analysis of high- and low-level deep learning approaches in microsatellite instability prediction"

**Supplementary Figure 1-3**

**Supplementary Table 1-6**

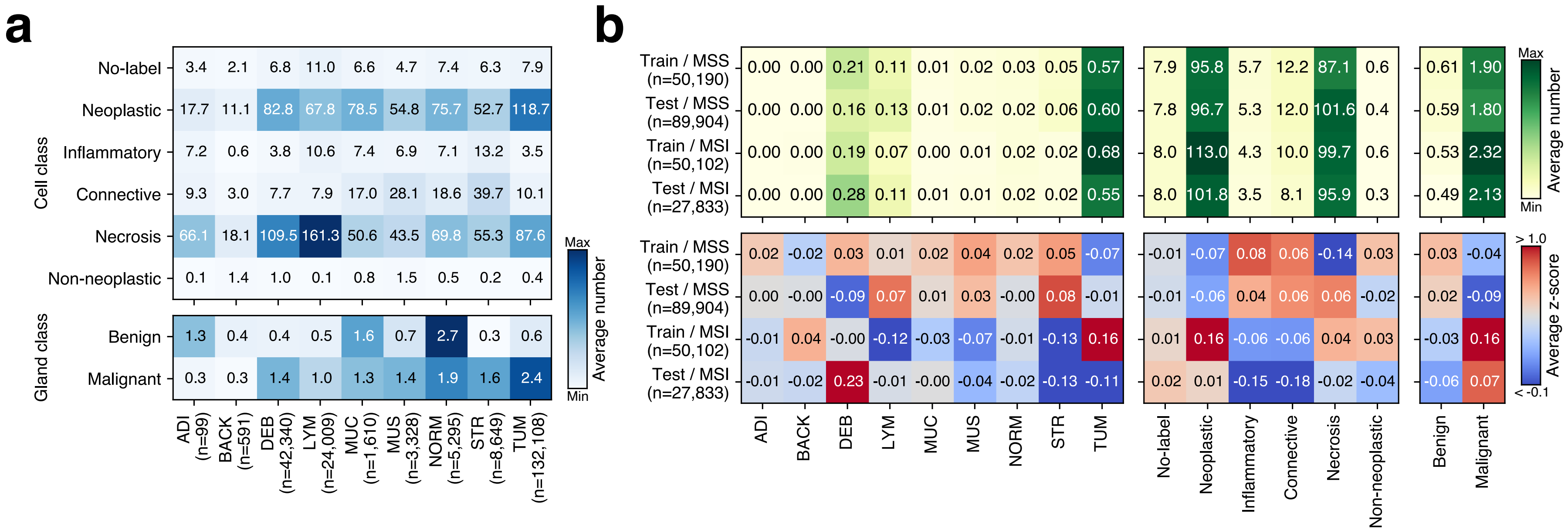


**Supplementary Figure 1.** (a) Correlations among low-level features derived from both the training and test sets from the stomach (STAD) cohort. (b) Low-level features of the training and test sets of the STAD cohort. Average number (top) and z score (bottom). ADI, adipose tissue; BACK, background; DEB, debris; LYM, lymphocytes; MUC, mucus; MUS, smooth muscle; NORM, normal colon mucosa; STR, cancer-associated stroma; TUM, colorectal adenocarcinoma epithelium. The heatmaps were created with Python v3.8.8 (https://www.python.org) and Matplotlib v3.4.3 (https://www.matplotlib.org). The figure was generated using Inkscape v1.1 (https://inkscape.org).

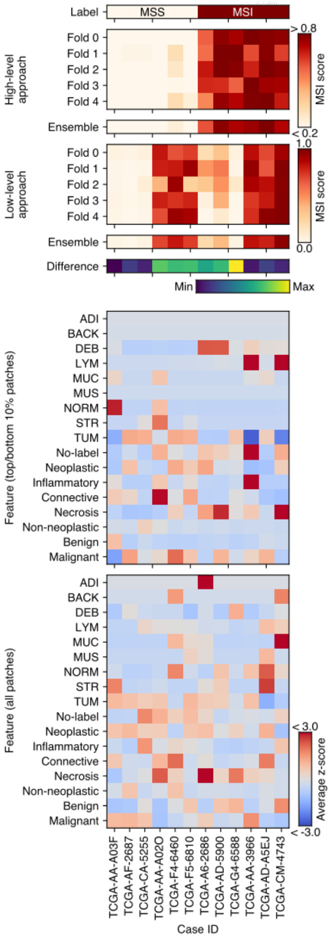


**Supplementary Figure 2.** Comparison of high/low-level approaches for select cases. Case-level prediction of EfficientNet (high-level approach) and LGBM (l ow-level approach) with MSI labels (top six panels). The predicted MSI scores of fivefold models and their ensemble are shown. Their difference is calculated based on the z scores of each ensemble. Representative image features derived from high/low-level approaches (bottom two panels). The average z score of the number of tissues, cells, and glands of the top 10\% of MSS/MSI patches and the case-level average. TCGA case IDs are shown as column labels. LGBM, lig ht gradient boosting machine; MSS, microsatellite stable; MSI, microsatellite instable (instability); TCGA, the cancer genome atlas; ADI, adipose tissue; BACK, background; DEB, debris; LYM, lymphocytes; MUC, mucus; MUS, smooth muscle; NORM, normal colon mucosa; STR, cancer-associated stroma; TUM, colorec tal adenocarcinoma epithelium. The heatmaps were created with Python v3.8.8 (https://www.python.org) and Matplotlib v3.4.3 (https://www.matplotlib.org). The fi gure was generated using Inkscape v1.1 (https://inkscape.org).

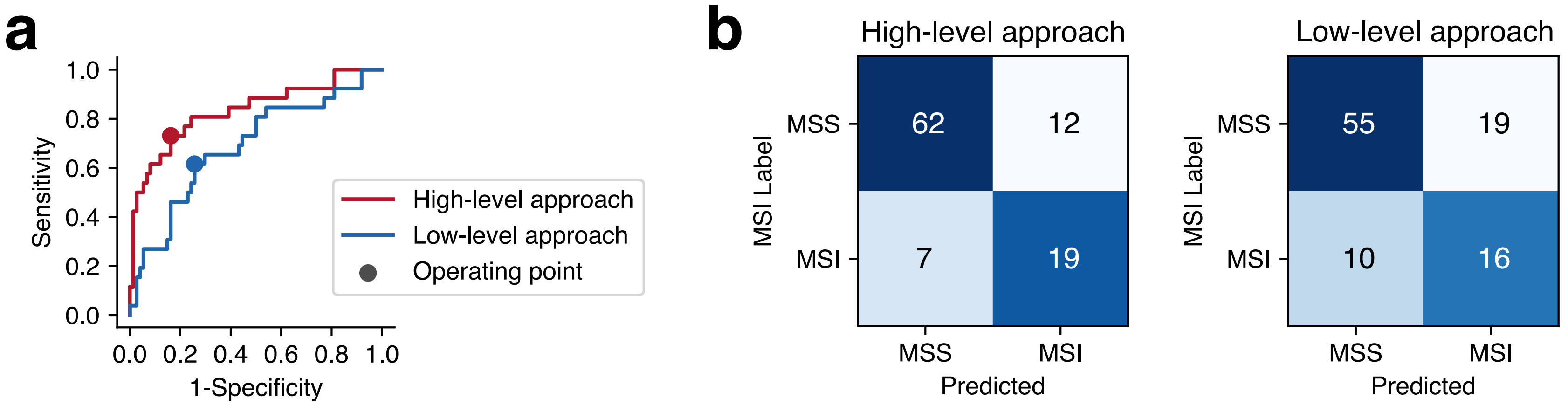


c

| Approach | AUROC | Threshold (operating point) | Accuracy | Balanced accuracy | Sensitivity (Recall) | Specificity | Precision | F1-score |
|---|---|---|---|---|---|---|---|---|
| High-level (ensembled) | 0.8316 | 0.40 | 0.8100 | 0.7843 | 0.7308 | 0.8378 | 0.6129 | 0.6667 |
| Low-level (ensembled) | 0.6866 | 0.35 | 0.7100 | 0.6793 | 0.6154 | 0.7432 | 0.4571 | 0.5246 |

**Supplementary Figure 3.** Classification performance of high-/low-level approaches. (a) Receiver operating characteristic curves of the high-level approach (red ) and low-level approach (blue) on the test set. For each approach, we used the probabilities of the ensemble of fivefold models. The operating point was dete rmined as the threshold at the point where the F1-score of the test set was the maximum, while the threshold was adjusted from 0 to 1 at 0.05 intervals. (b) Confusion matrix of each approach at the operating point. (c) Performance metrics of each approach. AUROC, area under the receiver operating characteristic c urve. The heatmaps were created with Python v3.8.8 (https://www.python.org) and Matplotlib v3.4.3 (https://www.matplotlib.org). The figure was generated using Inkscape v1.1 (https://inkscape.org).

| | Kather et al. | Ours |
|---|---|---|
| Implementation | Matlab | Pytorch |
| Architecture | ResNet-18 | [ResNet-18 v2, ShuffleNet v2 x1.0, EfficientNet-b0, ResNext-50] |
| Training | 12.5% held-out validation set | 5-fold cross validation |
| Early stopping tolerance | Three validation check | Six validation check |
| Validation check | 256 iterations | 256 iterations |
| Pretraining | ImageNet | ImageNet |
| Trainable layer | last 10 layers | [fully trainable, partial freeze] |
| Optimizer | Adam | Adam |
| L2-regularization | 1e-4 | 1e-4 |
| Leaning rate | 1e-6 | [1e-3, 1e-4, 1e-5, 1e-6] |
| Batch size | 256 | 256 |
| Max training duration | 100 epochs | 25,600 iteration |
| Augmentation | Geometric only | Geometric only |
| Confidence interval | 500-fold bootstrapping | 5 models from cross-validation |

**Supplementary Table 1. Comparison of a previous method and the search space used in this study.**

|  |  |  | No. of patches (No. of cases) |  |
|---|---|---|---|---|
| Cohort | Train/test | Fold # | MSS | MSI |
| CRC | Train set | 1 | 9,084 (45) | 9,781 (8) |
|  |  | 2 | 9,495 (44) | 9,187 (8) |
|  |  | 3 | 8,691 (44) | 11,347 (8) |
|  |  | 4 | 11,208 (44) | 7,750 (8) |
|  |  | 5 | 8,226 (44) | 8,639 (7) |
|  |  | 1+2+3+4+5 | 46,704 (221) | 46,704 (39) |
|  | Test set | - | 70,569 (74) | 28,335 (26) |
| STAD | Train set | 1 | 9,201 (30) | 11,858 (7) |
|  |  | 2 | 10,510 (30) | 9,988 (7) |
|  |  | 3 | 10,579 (30) | 10,497 (7) |
|  |  | 4 | 9,456 (30) | 7,884 (7) |
|  |  | 5 | 10,539 (30) | 10,058 (7) |
|  |  | 1+2+3+4+5 | 50,285 (150) | 50,285 (35) |
|  | Test set | - | 90,104 (74) | 27,904 (25) |

MSS, Microsatellite stable; MSI, Microsatellite unstable

**Supplementary Table 2. MSI dataset configuration**

| Architecture | Batch size | Learning rate | Total tensors | Trainable tensors | Trainable parameters | AUROC (95% CI) |
|---|---|---|---|---|---|---|
| ResNet-18 v2 | 256 | 1e-3 | 62 | 62 | 11177538 | 0.6967 (0.6551-0.7382) |
| ResNet-18 v2 | 256 | 1e-4 | 62 | 62 | 11177538 | 0.7639 (0.7242-0.8037) |
| ResNet-18 v2 | 256 | 1e-5 | 62 | 62 | 11177538 | 0.7833 (0.7306-0.8359) |
| ResNet-18 v2 | 256 | 1e-6 | 62 | 62 | 11177538 | 0.7693 (0.7345-0.8041) |
| ResNet-18 v2 | 256 | 1e-3 | 62 | 17 | 8394754 | 0.7816 (0.7420-0.8212) |
| ResNet-18 v2 | 256 | 1e-4 | 62 | 17 | 8394754 | 0.7933 (0.7473-0.8394) |
| ResNet-18 v2 | 256 | 1e-5 | 62 | 17 | 8394754 | 0.7991 (0.7720-0.8261) |
| ResNet-18 v2 | 256 | 1e-6 | 62 | 17 | 8394754 | 0.7938 (0.7635-0.8240) |
| ShuffleNet v2 x1.0 | 256 | 1e-3 | 170 | 170 | 1255654 | 0.7544 (0.7039-0.8049) |
| ShuffleNet v2 x1.0 | 256 | 1e-4 | 170 | 170 | 1255654 | 0.7433 (0.7068-0.7798) |
| ShuffleNet v2 x1.0 | 256 | 1e-5 | 170 | 170 | 1255654 | 0.7323 (0.7015-0.7632) |
| ShuffleNet v2 x1.0 | 256 | 1e-6 | 170 | 170 | 1255654 | 0.6243 (0.5571-0.6916) |
| ShuffleNet v2 x1.0 | 256 | 1e-3 | 170 | 47 | 980586 | 0.7504 (0.6957-0.8051) |
| ShuffleNet v2 x1.0 | 256 | 1e-4 | 170 | 47 | 980586 | 0.7451 (0.6975-0.7927) |
| ShuffleNet v2 x1.0 | 256 | 1e-5 | 170 | 47 | 980586 | 0.7280 (0.6800-0.7759) |
| ShuffleNet v2 x1.0 | 256 | 1e-6 | 170 | 47 | 980586 | 0.6047 (0.4952-0.7141) |
| EfficientNet-b0 | 96 | 1e-3 | 213 | 213 | 4010110 | 0.7395 (0.6689-0.8101) |
| EfficientNet-b0 | 96 | 1e-4 | 213 | 213 | 4010110 | 0.7983 (0.7606-0.8361) |
| EfficientNet-b0 | 96 | 1e-5 | 213 | 213 | 4010110 | 0.7917 (0.7734-0.8100) |
| EfficientNet-b0 | 96 | 1e-6 | 213 | 213 | 4010110 | 0.7673 (0.7250-0.8095) |
| EfficientNet-b0 | 96 | 1e-3 | 213 | 41 | 1895698 | 0.7946 (0.7263-0.8629) |
| EfficientNet-b0 | 96 | 1e-4 | 213 | 41 | 1895698 | 0.8052 (0.7602-0.8501) |
| EfficientNet-b0 | 96 | 1e-5 | 213 | 41 | 1895698 | **0.8065 (0.7758-0.8373)** |
| EfficientNet-b0 | 96 | 1e-6 | 213 | 41 | 1895698 | 0.7267 (0.6765-0.7770) |
| Resnext50 | 84 | 1e-3 | 161 | 161 | 22984002 | 0.6646 (0.5398-0.7893) |
| Resnext50 | 84 | 1e-4 | 161 | 161 | 22984002 | 0.7968 (0.7696-0.8240) |
| Resnext50 | 84 | 1e-5 | 161 | 161 | 22984002 | 0.7819 (0.7543-0.8096) |
| Resnext50 | 84 | 1e-6 | 161 | 161 | 22984002 | 0.7871 (0.7790-0.7953) |
| Resnext50 | 84 | 1e-3 | 161 | 32 | 14548994 | 0.7825 (0.7279-0.8372) |
| Resnext50 | 84 | 1e-4 | 161 | 32 | 14548994 | 0.7640 (0.6825-0.8455) |
| Resnext50 | 84 | 1e-5 | 161 | 32 | 14548994 | 0.7741 (0.7482-0.8000) |
| Resnext50 | 84 | 1e-6 | 161 | 32 | 14548994 | 0.7870 (0.7637-0.8103) |

**Supplementary Table 3. Optimization of the MSI classification model**

| Architecture | Batch size | Learning rate | Total tensors | Trainable tensors | Trainable parameter | Average Accuracy (95%CI) | Average balanced accuracy (95%CI) |
|---|---|---|---|---|---|---|---|
| EfficientNet-b0 | 96 | 1e-3 | 213 | 213 | 4010110 | 0.7434 (0.6721-0.8147) | 0.7245 (0.6366-0.8124) |
| EfficientNet-b0 | 96 | 1e-4 | 213 | 213 | 4010110 | 0.9409 (0.9206-0.9613) | 0.9296 (0.9167-0.9425) |
| EfficientNet-b0 | 96 | 1e-5 | 213 | 213 | 4010110 | **0.9523 (0.9424-0.9623)** | **0.9314 (0.9160-0.9468)** |
| EfficientNet-b0 | 96 | 1e-6 | 213 | 213 | 4010110 | 0.9460 (0.9419-0.9501) | 0.9202 (0.9121-0.9284) |
| EfficientNet-b0 | 96 | 1e-3 | 213 | 41 | 1895698 | 0.9283 (0.9188-0.9379) | 0.8949 (0.8813-0.9086) |
| EfficientNet-b0 | 96 | 1e-4 | 213 | 41 | 1895698 | 0.9379 (0.9358-0.9400) | 0.9084 (0.9029-0.9140) |
| EfficientNet-b0 | 96 | 1e-5 | 213 | 41 | 1895698 | 0.9380 (0.9359-0.9401) | 0.9078 (0.9043-0.9113) |
| EfficientNet-b0 | 96 | 1e-6 | 213 | 41 | 1895698 | 0.9379 (0.9358-0.9400) | 0.9084 (0.9029-0.9140) |

**Supplementary Table 4. Optimization of the tissue classification model**

| | F1-score | |
|---|---|---|
| | Test A | Test B |
| All | 0.9019 | 0.7290 |
| Benign | 0.9213 | 0.9143 |
| Malignant | 0.8488 | 0.6927 |

**Supplementary Table 5. Evaluation of the gland detection model on test sets of the GlaS dataset**

| Lambda | Learning rate | Feature fraction | AUROC (95% CI) |
|---|---|---|---|
| 0 | 0.001 | 0.5 | 0.6679 (0.6219-0.7139) |
| 0 | 0.001 | 0.8 | 0.6658 (0.6185-0.7131) |
| 0 | 0.001 | 1 | 0.6633 (0.6292-0.6973) |
| 0 | 0.01 | 0.5 | 0.6656 (0.6194-0.7118) |
| 0 | 0.01 | 0.8 | 0.6677 (0.6219-0.7134) |
| 0 | 0.01 | 1 | 0.6690 (0.6235-0.7146) |
| 0 | 0.1 | 0.5 | 0.6501 (0.6095-0.6907) |
| 0 | 0.1 | 0.8 | 0.6634 (0.6249-0.7020) |
| 0 | 0.1 | 1 | 0.6659 (0.6275-0.7043) |
| 0.1 | 0.001 | 0.5 | 0.6695 (0.6242-0.7149) |
| 0.1 | 0.001 | 0.8 | 0.6634 (0.6241-0.7027) |
| 0.1 | 0.001 | 1 | 0.6657 (0.6317-0.6996) |
| 0.1 | 0.01 | 0.5 | 0.6674 (0.6220-0.7127) |
| 0.1 | 0.01 | 0.8 | 0.6709 (0.6235-0.7183) |
| 0.1 | 0.01 | 1 | 0.6748 (0.6251-0.7246) |
| 0.1 | 0.1 | 0.5 | 0.6544 (0.6085-0.7002) |
| 0.1 | 0.1 | 0.8 | **0.6749 (0.6365-0.7134)** |
| 0.1 | 0.1 | 1 | 0.6629 (0.6333-0.6925) |
| 0.2 | 0.001 | 0.5 | 0.6662 (0.6245-0.7079) |
| 0.2 | 0.001 | 0.8 | 0.6632 (0.6239-0.7025) |
| 0.2 | 0.001 | 1 | 0.6544 (0.6180-0.6907) |
| 0.2 | 0.01 | 0.5 | 0.6644 (0.6188-0.7101) |
| 0.2 | 0.01 | 0.8 | 0.6699 (0.6242-0.7155) |
| 0.2 | 0.01 | 1 | 0.6684 (0.6209-0.7159) |
| 0.2 | 0.1 | 0.5 | 0.6570 (0.6161-0.6978) |
| 0.2 | 0.1 | 0.8 | 0.6705 (0.6424-0.6985) |
| 0.2 | 0.1 | 1 | 0.6599 (0.6197-0.7001) |

**Supplementary Table 6. Optimization of the low-level approach**
